# The synergistic anti-Warburg efficacy of temozolomide, metformin and epigallocatechin gallate in glioblastoma

**DOI:** 10.1101/2022.11.22.517597

**Authors:** Shreyas S. Kuduvalli, Daisy S Precilla, Indrani Biswas, T. S Anitha

## Abstract

**Background:** An important hallmark of glioblastoma aggressiveness is its altered metabolism of glucose. This metabolic shift wherein the tumor cells employ aerobic glycolysis regardless of oxygen availability via reprogramming of mitochondrial oxidative phosphorylation is known as the Warburg effect. Previous literatures have linked this metabolic reprograming to tumor progression glioblastoma cell proliferation making it a key target for targeted drug therapy.

**Objective:** To evaluate the anti-Warburg efficacies of the triple-drug combination of temozolomide, metformin and epigallocatechin gallate in preclinical glioblastoma models.

**Methodology:** Based on this lacuna, the current study aimed to explore the therapeutic efficacy of the triple-drug combination of temozolomide, metformin and epigallocatechin gallate in attenuating Warburg effect and glucose uptake in glioblastoma both *in vitro* and *in vivo*.

**Results:** Our results showed that the triple-drug combination had significantly reduced glucose uptake and reversed the Warburg effect in glioblastoma cells and in the xenograft-induced glioma rat model.

**Conclusion:** Thus, the triple-drug combination would serve as an effective therapeutic regime to hamper glioblastoma progression via altering glucose metabolism and improve the overall prognosis in patient setting.

## 1. Introduction

Cellular homeostasis is mainly dependent on the energy derived by the metabolism of nutrients from the surrounding environment [1]. In our body among all the organs, brain has very high metabolism accounting for approximately 20% and 60% of the body’s total oxygen and glucose uptake respectively. In addition, as brain lacks the ability to store glucose as glycogen, there exists a need for constant supply of glucose, thereby making glucose as one of the most abundantly available macromolecule in the brain [2,3]. With regard to the metabolism of glucose in the brain, it is well-known that the neurons and glial cells employ tricarboxylic acid (TCA) cycle to generate ATP [3]. In contrast, during neoplastic state, GB and other glioma cells preferentially metabolize glucose to lactate even in the presence of ample oxygen which is known as the “Warburg” effect. This metabolic shift has been reported to enable tumor cells generate sufficient ATP by metabolising glucose independent of oxygen levels [1,4].

Glioblastoma (GB), the most invasive type of glioma developed from astrocytes, is one of the most lethal primary tumors of the central nervous system (CNS). The median survival for patients with GB remains under 14 to 15 months, following standard-of-care therapy with surgery, radiation and chemotherapy [5]. The only standard-drug for primary GB is Temozolomide (T) that has shown to exhibit clinically proven effects; however, the advantages are not palliative due to the development of chemo-resistance against the standard drug by the GB cells [6]. In addition to these aggressive hallmarks such as acquired chemo-resistance, sustaining proliferation and deregulated glucose metabolism pathways contribute to the poor prognosis for GB [7]. This metabolic reprograming has been marked as an important accomplice that directly contributes to GB tumorigenesis as well as enhanced chemoresistance, which makes it a novel target in GB therapy [8].

GB cells are known to divert the carbon derived from glucose metabolism into the pentose phosphate pathway (PPP) to combat oxidative stress. The oxidative PPP employs glucose-6-phosphate as substrate to generate NADPH. This non-oxidative reactions of PPP lead to the production of ribose-5-phosphate (R5P) that is mainly involved in nucleotide biosynthesis [9]. Furthermore, glycolytic enzymes such as phosphofructokinase 1 (PFKP) and pyruvate kinase M2 (PKM2) are highly regulated in GB, which results in the over-expression of transketolases that enhances tumor growth and proliferation in various cancers including GB [9–12]. To encounter this high metabolic demand of glucose, glucose transport (GLUT) proteins such as GLUT1, and to a lesser extent GLUT3 and GLUT4, have been reported to be increased in GB cells. Additionally, signalling pathways, upstream of this metabolic shift leading to tumor invasiveness have been linked to increased levels of glucose transporters in GB [10,13]. This reprograming of glucose metabolism that initiates with glycolysis leads to the formation of pyruvate by activating pyruvate kinase M2 (PKM2), which is also the final step of glycolysis in normal and cancer cells [14]. Contrary to normal cells, the isoenzymes, lactate dehydrogenase V (LDHV) catalyses pyruvate to lactate instead of oxidative phosphorylation via citric acid cycle in cancer cells [15]. However, this excessive non-oxidative metabolic breakdown of glucose leads to the accumulation of monocarboxylic acids such as lactate, that plays a major role in GB homeostasis [16]. In response to counteract this accumulation of lactate, cells produce monocarboxylate transporter 1 & 4 (MCT1 & 4) which to our current understanding, provides a key role in the import and export of lactate, respectively. While the major function of MCT4 is to export lactate, MCT1 has been mutually characterized to maintain the intracellular influx of lactate [17].

In our pervious study, we employed the drugs, T, metformin (M) and epigallocatechin gallate (E) to study their anti-GB efficacies both individually and in combination in both *in vitro* and *in vivo* orthotopic xenograft experimental models [18] wherein, the triple-drug combination of TME effectively attenuated GB via oxidative stress-mediated inactivation of PI3K/AKT/mTOR leading to apoptosis. To further gain insight into the effect of these drugs on glucose metabolism in GB, this study aimed to analyse the efficacy of our combination of drugs in reversing Warburg effect in GB cells by analysing the levels of GLUT1 & 4, PKM, LDHV and MCT 1 & 4.

## 2. Methodology

### 2.1 Cell line and culture conditions

Human GB cell line, U-87 MG and rat glioma cell line C6 were obtained from National Centre for Cell Science (NCCS), Pune, India. The cells were sub-cultured in Dulbecco’s Modified Eagle’s Medium (DMEM) supplemented with 10% FBS at 5% CO_2_ and 37°C. On reaching 85% confluence, the cells were harvested using 0.25% trypsin and used accordingly. For the treatment, the stock solution (10 mM) of each drug was made with their respective solvents and were further diluted using serum-free medium to arrive at the required concentrations as in our previous study [18]. Cells that were treated with vehicle alone were considered as control.

### 2.2 Xenograft Glioblastoma model

Healthy male Wistar rats (180–240g / 6 to 8 weeks old) were purchased from college of veterinary and animal sciences, Mannuthy, Kerala and were maintained in accordance with institutional guidelines and regulations of Sri Balaji Vidyapeeth (Deemed to-be University), Puducherry, India. All the animal experimental protocols were reviewed, approved and performed in accordance with the guidelines and regulations set by the Institutional Animal Ethics Committee (IAEC) Mahatma Gandhi Medical College and Research Institute, Sri Balaji Vidyapeeth (Deemed to-be University), Puducherry (07/IAEC/MG/08/2019-II). All experimental procedures were performed in accordance with COPE guidelines.

Wistar rats were anesthetized with a ketamine/xylazine cocktail solution (87 mg/kg body weight and 13 mg/kg body weight) and were placed in the stereotaxic head frame. A 1 cm midline scalp incision was made, and 1×10^6^ C6 glioma cells in 3 μl phosphate buffer saline (PBS) was injected at a depth of 6.0 mm in the right striatum (coordinates with regard to Bregma: 0.5 mm posterior and 3.0 mm lateral) through a burr hole in the skull using a 10-μl Hamilton syringe to deliver tumor cells to a 3.5-mm intra-parenchymal depth. The burr hole in the skull was sealed with bone wax and the incision was closed using dermabond. The rats were monitored daily for signs of distress and death. The study design and treatment groups are given in table 1.

**Table 1:** Drug dosage and its formulation

| Groups | Drugs | No. of animals (48) | Dosage<sup>#</sup> (mg/Kg body weight) |
| --- | --- | --- | --- |
| Group I | C | 03 | NA |
| Group II | TC | 03 | NA |
| Group III | T | 06 | T (10) |
| Group IV | M | 06 | M (15) |
| Group V | E | 06 | E (25) |
| Group VI | TM | 06 | T+M (10+15) |
| Group VII | TE | 06 | T+E (10+25) |
| Group VIII | ME | 06 | M+E (15+25) |
| Group IX | TME | 06 | T+M+E (10+15+25) |

**Table 2.**
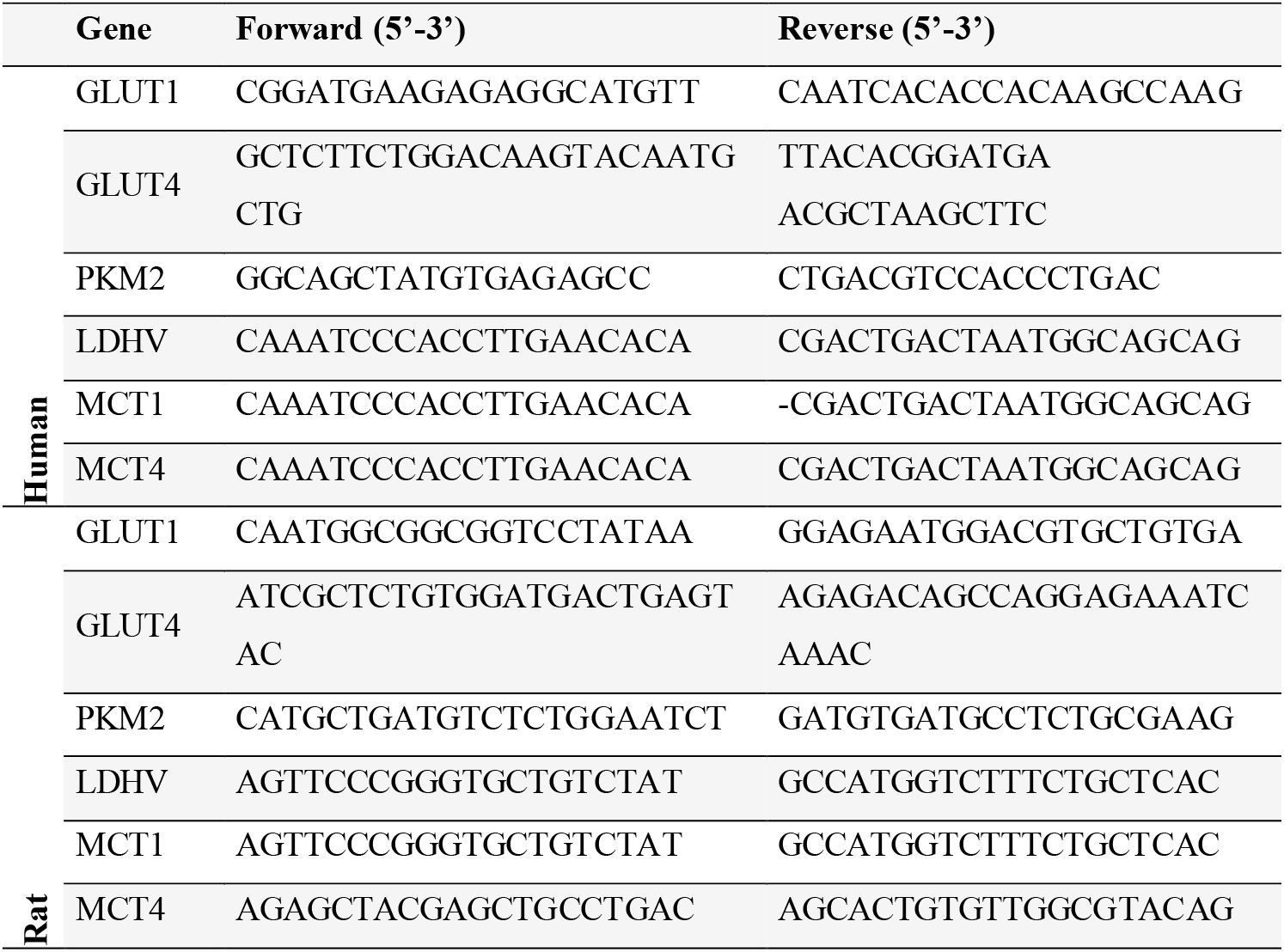
list of primers employed in the study

### 2.3 Primary culture of astrocytes

Primary astrocytes were collected from the isolated brain tissues as described by Schildge, *et al*. [19]. Briefly, the brain tissues of all the experimental groups were isolated, immediately after decapitation. The minced tissue was dissociated using 0.05% trypsin at 37^0^C for 30 min. To isolate individual astrocytes, the cell suspension was passed through 40 μm strainer. At last, astrocytes were seeded at a density of 1.5×10^5^ cells/cm^2^ in 90% DMEM containing 10% FBS, and 4.5 g/L glucose. The astrocytes were cultured at 37^0^C with 95% air and 5% CO_2_.

### 2.4 Glucose uptake assay

Glucose uptake assay was performed using Glucose Uptake-Glo kit (Promega, USA). The assay was performed according to the manufacturer’s protocol, for both the cell lines and astrocytes isolated from the rat brain tissue.

### 2.5 Quantitative real-time polymerase chain reaction (qRT-PCR)

The total RNA was extracted from the cell lysates and brain tissues of all the experimental groups using TRIZOL reagent (Takara Bio, Shiga, Japan). Quantitative RT-PCR was carried out using a CFX96 thermo cycler (Bio-Rad, California, USA) and TB Green Premix Ex Taq I (Takara Bio, Shiga, Japan) to detect messenger ribonucleic acid (mRNA). The specific PCR primer sequences are listed in Supplementary Table 2. Independent experiments were conducted in triplicate. The relative changes in gene expression were calculated with the 2^-ΔΔCt^ method, where ΔΔCt = (Ct target gene – Ctβ-actin) sample - (Sample Ct target gene – Control Ct target gene) calibrator.

### 2.6 Enzyme-Linked Immune Sorbent Assay

ELISA for U-87 MG, C6 cells and tumor-induced brain tissues was performed to quantify the protein levels GLUT-1 (Human: MBS8800801; Rat: MBS8805788), PKM2 (Human: MBS8800515; Rat: MBS8800516), LDHV (Human: MBS8801237; Rat: MBS8804669) and MCT-1 (Human: MBS8803496; Rat: MBS8806122) (My BioSource, California, USA). The concentration levels of the above-mentioned proteins were calculated using a standard curve obtained.

### 2.7 Statistical analysis

All the experiments were performed at least thrice. All quantitative data values are expressed as the mean ± standard deviation (SD). All the experimental data was analysed using Two-way Analysis of Variance (ANOVA) by GraphPad software (California, USA). p<0.01 was statistically significant.

## 3. Result and Discussion

### 3.1 The triple-drug combination hindered the uptake of glucose

Most cancer including GB are characterised by enhanced metabolic rates often accompanied by a high uptake of glucose [20]. This high level of glucose enhances GB progression and invasiveness. Consequently reduction in the uptake of glucose has been reported to hinder proliferation, apoptosis and enhanced survival in GB [16]. Based on these lacunae, we analysed the efficacy of the chosen drugs on glucose uptake levels in U-87 MG and C6 glioma cells as well as in rat brain tissue to determine if the chosen drugs either individually or as a combination could contribute to reduction in the uptake of glucose levels. Interestingly, we observed that the triple-drug treatment (TME) showed a significant reduction in glucose uptake both *in vitro* and *in vivo* (p<0.001) (**Fig.1**). Further, the dual treatment of TM and individual treatment with M had also reduced glucose uptake significantly in both the cell lines and in the brain tissues of GB-bearing xenograft rats (p<0.01 & p<0.05) (**Fig.1**). This was in accords with previous studies which have reported the ability of T and E to individually reduce the uptake of glucose in various solid tumors including GB [21,22]. One possible reason for this significant reduction in glucose uptake might be attributed to reduction in the expression of glucose transporters, GLUT1, GLUT4, PKM and LDHV, which has been reported to play a crucial role in controlling the demand for glucose. From this, it was evident that treatment with TME in this study significantly reduced the uptake of glucose in GB cells.

**Fig. 1.**
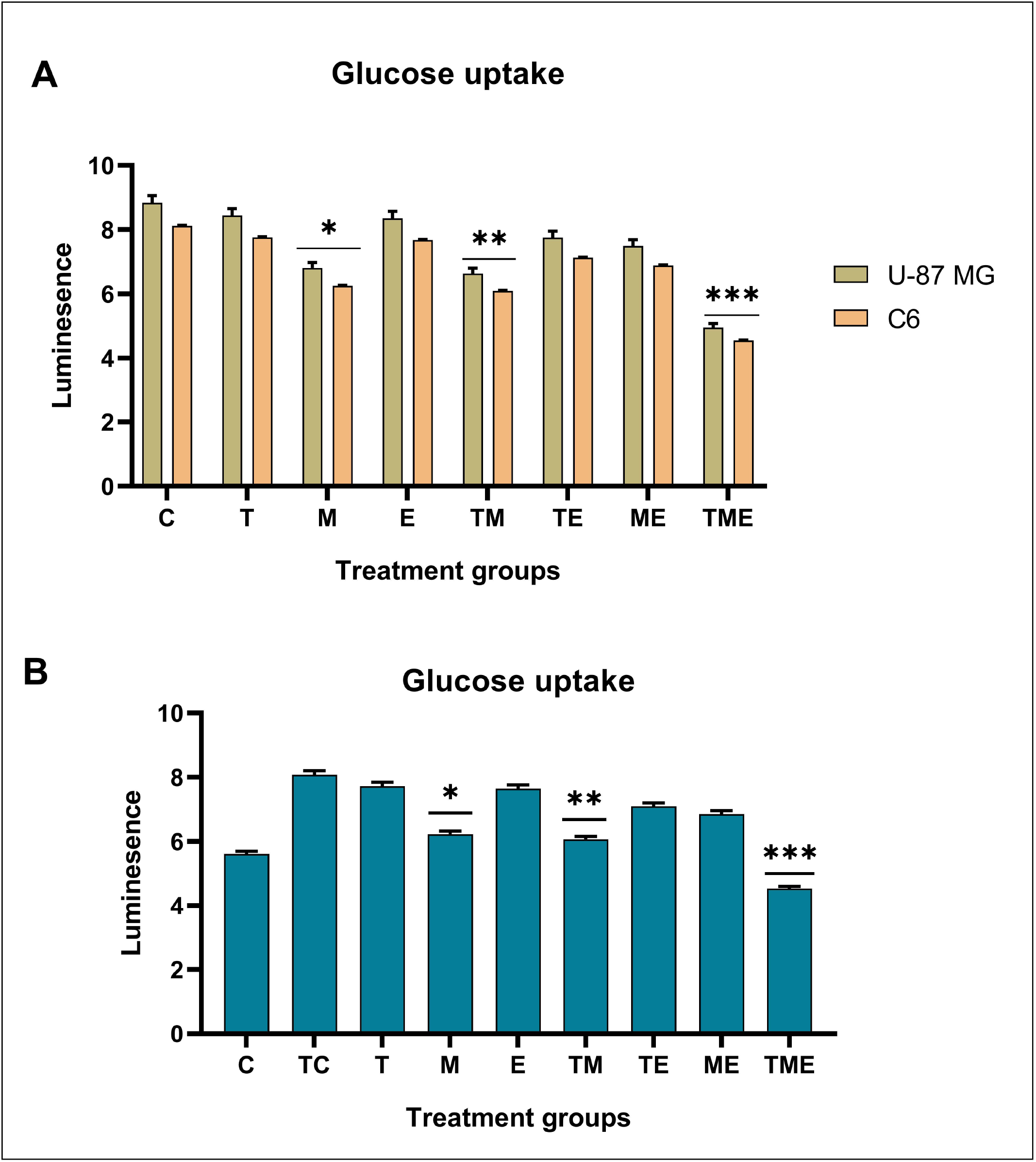
Glucose uptake by the GB cells and isolated astrocytes in various treatment groups. (**A**) U-87MG & C6 GB cells (**B**) astrocytes isolated from the tumor tissue (*p<0.05, **p<0.01, ***<0.001)

### 3.2 The anti-Warburg effect of the triple-drug combination

Previous studies have reported that most solid tumors such as GB are often characterised by a dense hypoxic core, with depleted levels of oxygen [23]. To overcome this hypoxic effect, cells of the GB tumor facultatively opt for the non-oxidative mode of glucose metabolism. Also, studies have symbolised that, Warburg effect with a high intake of glucose is facilitated by the over expression of the glucose transporter proteins such as GLUT1 and GLUT4. Glucose is initially metabolised to pyruvate by the action of PKM2 [24,25]. Under normal conditions, this pyruvate is delocalised to the mitochondria and is followed by citric acid cycle (oxidative phosphorylation) to generate ATP. However, in cancers cells, lactate dehydrogenase V (LDHV), an isoenzyme which possess a high affinity to pyruvate, catalyses the conversion of pyruvate to lactate, acting as a key regulator of this metabolic switch. This results in the accumulation of abundant amount of lactate which is then regulated throughout the tumor with the help of monocarboxylate transporter proteins (MCT 1 & 4). The efficient regulation of inter and intra-cellular levels of lactate has also been reported to facilitate tumor cell proliferation and GB progression.

Following this concept, we analysed the gene and protein expression levels of key markers of Warburg effect (GLUT 1&4, PKM2, LDHV, MCT 1&4). Interestingly, our expression studies highlighted that the gene expression and protein levels of theses markers were highly upregulated in the non-treated GB cells and TC group, whereas, the triple dug combination (TME) had significantly reduced the levels of all these markers in both the glioma cell lines as well as in GB-tissue bearing xenografts (**p<0.001**; **Fig.2**). Also, the dual-drug treatment of TM too had significantly reduced the levels of these markers in both GB cells and xenografts (**p<0.01**; **Fig.2**). Whilst the other treatment groups did not show any significant effects. This indicated that the triple-drug combination of TME effectively inhibited Warburg effect in GB cells both *in vitro* and *in vivo*.

**Fig. 2.**
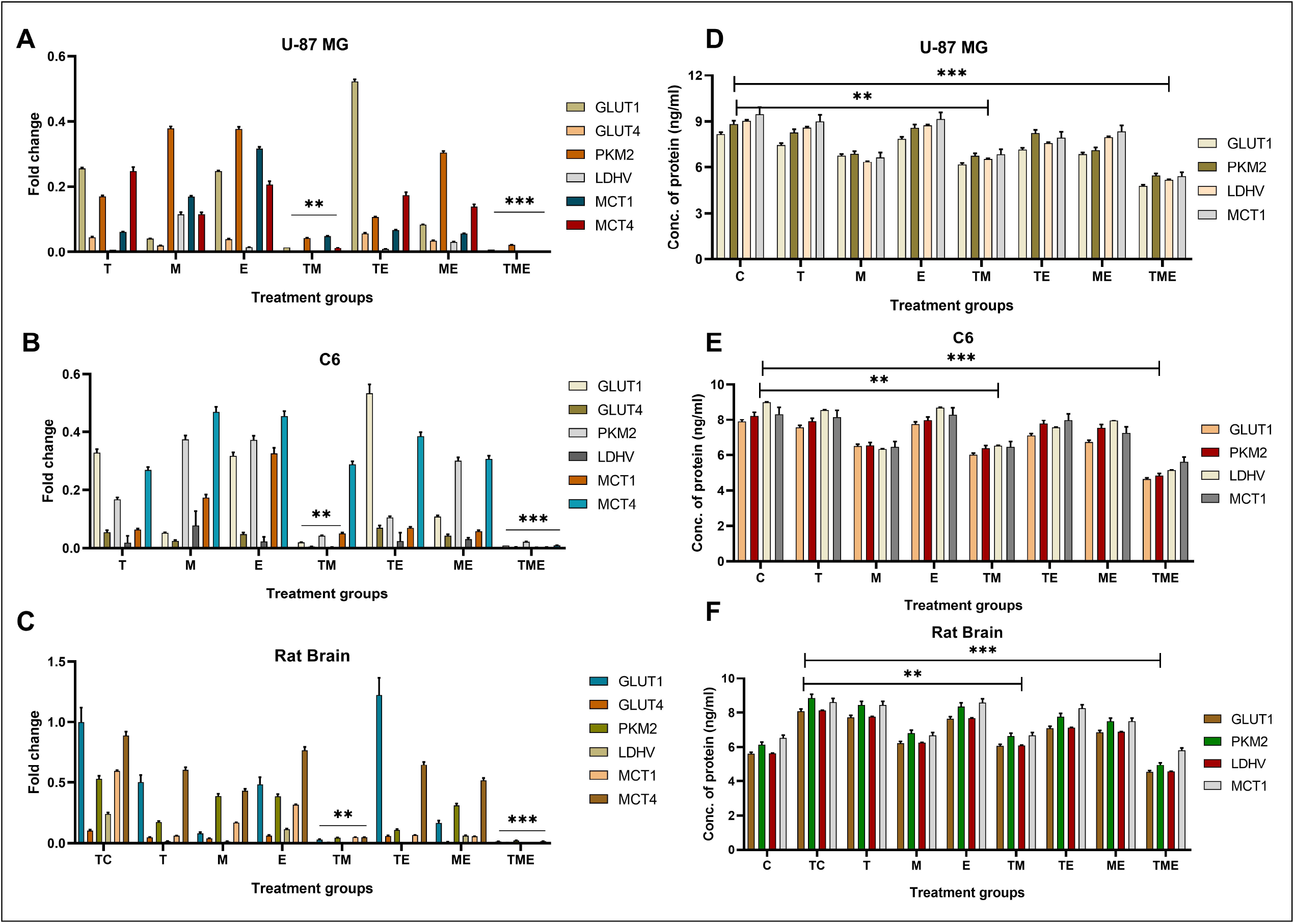
**(A-C)** Gene expression levels of GLUT1 & 4, PKM2, LDHV and MTC1 & 4. (A) U-87 MG Human GB cell line, (B) C6 rat GB cell line and (C) Rat brain tissue. The triple-drug combination most significantly supressed the gene expression of all the makers of Warburg effect followed by the dual treatment with TM. **(D-F)** Protein levels of GLUT1, PKM2, LDHV and MTC1. (D) U-87 MG Human GB cell line, (E) C6 rat GB cell line and (F) Rat brain tissue. The triple-drug combination of TME followed by dual combination of TM significantly protein levels GLUT1, PKM2, LDHV and MCT1 (**p<0.01, ***<0.001)

## 4. Conclusion

Taken together, the results of the current study highlighted that the triple-drug combination of TME mechanistically inhibited glucose uptake and attenuated Warburg effect both *in vitro* and *in vivo*, suggesting that this combination can be used as a targeted drug therapy to attenuate GB. However, further studies in this regard are warranted to conclusively determine downstream effects of this triple-drug combination following modulations of glucose metabolic pathway.

## 6. Availability of data and materials

Not applicable.

## 8. Ethics approval and consent to participate

Not applicable.

## 9. Human and animal rights

All the animal experimental protocols were reviewed, approved and performed in accordance with the guidelines and regulations set by the Institutional Animal Ethics Committee (IAEC) Mahatma Gandhi Medical College and Research Institute, Sri Balaji Vidyapeeth (Deemed to-be University), Puducherry (07/IAEC/MG/08/2019-II).

## 10. Consent for publication

Not applicable.

## 11. List of abbreviations

GB: Glioblastoma Multiforme
T: Temozolomide
M: Metformin
E: Epigallocatechin Gallate
TME: Triple-drug combination of temozolomide, metformin and epigallocatechin gallate

## 13. Author contribution

Dr, Anitha T. S. has formulated and supervised the work and assisted in interpretation of results. Mr. Shreyas S. Kuduvalli performed all the practical procedures and analysed the results. Ms. Daisy Precilla and Ms. Indrani Biswas has helped in drafting the manuscript. All authors have read and approved the final manuscript.

## 14. Acknowledgement

We would like to thank Sri Balaji Vidyapeeth (Deemed to-be University) for providing the basic research facility and instrumentation.

